# PanGraphViewer: A Versatile Tool to Visualize Pangenome Graphs

**DOI:** 10.1101/2023.03.30.534931

**Authors:** Yuxuan Yuan, Ricky Ka-Kui Ma, Ting-Fung Chan

## Abstract

Pangenome graphs provide a powerful way to present both sequence and structural features in a given genome relative to the typical features of a population. There are different methods of building pangenome graphs, but few tools are available to visualize them. To address this problem, we developed PanGraphViewer, which is written in Python 3 and runs on all major operating systems. The PanGraphViewer package contains two separate versions: a desktop-based application and a web-based application. Compared to other graph viewers that are initially designed to visualize individual genome graphs, PanGraphViewer targets pangenome graphs and allows the viewing of pangenome graphs built from multiple genomes in either the (reference) graphical fragment assembly format or the variant call format (VCF). Apart from visualization of different types of structural variations (SV), PanGraphViewer also integrates genome annotations with graph nodes to analyze insertions or deletions in a particular gene model. The graph node shapes in PanGraphViewer can represent different types of genomic variations when a VCF file is used. Notably, PanGraphViewer displays subgraphs from a chromosome or sequence segment based on any given coordinates. This function is absent from most genome graph viewers. PanGraphViewer is freely available at https://github.com/TF-Chan-Lab/panGraphViewer to facilitate pangenome analysis.

## Introduction

With the large number of high-quality genome assemblies being produced, it is recognized that a single genome assembly cannot capture the entire genomic content of a species [1]. A single genome assembly may impede the understanding of genomic changes in individuals that may be associated with important phenotypic traits. The pangenomics concept was conceived to address this problem, and it has been applied to a variety of species, such as humans, wheat, barley, and soybean [2–6].

During the construction of a pangenome, although different approaches have been proposed [2], most rely on a linear genome assembly to capture the genetic features of a species, with some novel contigs that cannot be efficiently anchored to the main sequences. Practically, the commonly used pangenome construction approaches identify the most commonly shared and most variable sequences in the studied individuals. However, they cannot efficiently characterize genomic variations for instance, allelic or heterozygous variations in the studied samples, thereby preventing a full understanding of the pangenome. To resolve this problem, pangenome graph approaches have recently been suggested [7].

A pangenome graph differs from a linear genome, which can only show linear connections in a pangenome assembly. A pangenome graph uses nodes to represent sequence segments and different paths to represent variations. By constructing node connections, theoretically, a pangenome graph can capture all of the genomic features or variations in the studied species [8]. The nodes and paths can easily characterize the genomic information that is shared among or unique to particular individuals. By exploring the nodes and paths, users can intuitively understand the pangenome graph.

Pangenome graphs may be constructed using different tools, such as vg [9] and minigraph [10]. However, few tools are available to visualize these graphs, making it difficult to comprehensively explore them. Previously developed genome graph viewers, such as Bandage [11], GfaViz [12] and Gfaestus (https://github.com/chfi/gfaestus), may be used to visualize a pangenome graph. However, those tools mainly focus on displaying the overall structure of the graph [7] and there are some pitfalls in using them. For example, Bandage and GfaViz require large amounts of computational resources to load and render a pangenome graph with tens of thousands of nodes and connections. This makes the application of the two software difficult for users with limited computing resources. The installation of GfaViz and Gfaestus is complicated for users with a limited computer science background, because it is non-trivial to install and set up the dependencies. Additionally, users cannot directly use position coordinates to visualize specific pangenomic regions, unless they zoom in and out of the entire pangenome graph to locate the regions of interest or using a third-party tool to get the node ids first and then browse the underlying subgraph. Moreover, these tools do not provide a function to connect genome annotations with graph nodes. Taken together, those genome graph viewers cannot fully meet the current demands in visualizing a pangenome graph.

Although several tools have been specifically developed to assist pangenome graph visualization, for instance, Sequence Tube Map [13] and MoMI-G [14], there are still some disadvantages in using them. For instance, both of the tools rely on vg and the data formats accepted are from vg [9]. This design may prevent data sharing with other pangenome graph viewers that do not support the vg formats. In addition, they use a tube layout and this layout may impede the viewing of graph components in a large-scale. To resolve the problems mentioned above, we have developed PanGraphViewer, a free software package to assist pangenome graph visualization on different operating systems with the adoption of the reference graphical fragment assembly (rGFA) format, GFA_v1 format and the variant call format (VCF). The mainly used layout is the fast compound spring embedder (fcose) layout. Overall, PanGraphViewer could increase the flexibility of using a pangenome graph to visualize the genomic variations and assist pangenome studies.

## Methods

### Design strategy

During the design of PanGraphViewer, two aspects were mainly concerned: system requirements and visualisation capability. System requirements refer to the environments that can host the application, which involve the operating system, software dependences and hardware configurations, while visualisation capability refers to the components that can be visualised in a pangenome graph. To fit the design purposes, we have developed PanGraphViewer with two different versions targeting for users with different system environments and computer science backgrounds. To meet the pangenome graph visualization demand, we have enabled PanGraphViewer to accept different pangenome graph formats and allowed the visualization of subgraphs with different elements displayed, for example structural variations in a variation-based pangenome graph. To further assist users to visualize pangenomic regions of interest in an efficient way, in PanGraphViewer, we have prepared a function to allow users to specify coordinates to visualize the underlying subgraph without the assistance from a third-party tool.

### Tool versions

PanGraphViewer is written in Python 3 and runs on Microsoft Windows, Linux, and macOS operating systems. A desktop-based version and a web-based version are provided in the package. Basically, most functions in the two PanGraphViewer versions are the same; however, there are some variations (**Table 1**). For example, the desktop-based PanGraphViewer was designed using PyQt5, with a provision to further extend and update through an application programming interface, while the web-based version was created using Django3.1.3 to be a light web application. The desktop-based PanGraphViewer is targeted for a single user and optimized for performance, while the web-based PanGraphViewer is designed for multiple users and optimized for accessibility and portability. Additionally, for the web-based PanGraphViewer, installation only needs once in a server, and users could freely access the tool using a static IP and the user account. To ease the deployment for the web-based version, a docker image is also provided.

**Table 1.**
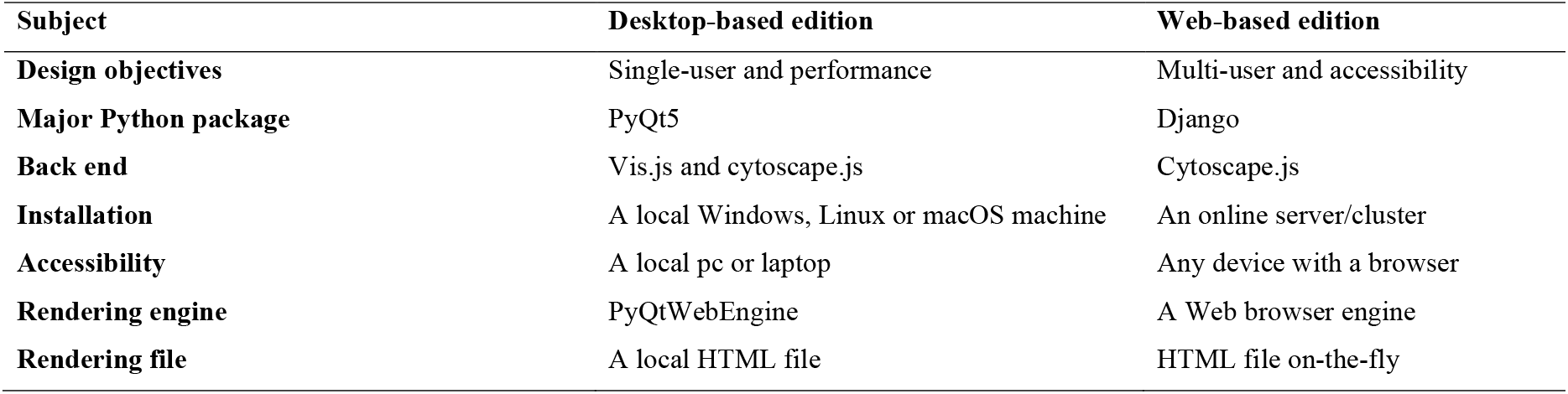
Major differences between the desktop-based PanGraphViewer and the web-based PanGraphViewer.

| **Subject** | **Desktop-based edition** | **Web-based edition** |
| --- | --- | --- |
| **Design objectives** | Single-user and performance | Multi-user and accessibility |
| **Major Python package** | PyQt5 | Django |
| **Back end** | Vis.js and cytoscape.js | Cytoscape.js |
| **Installation** | A local Windows, Linux or macOS machine | An online server/cluster |
| **Accessibility** | A local pc or laptop | Any device with a browser |
| **Rendering engine** | PyQtWebEngine | A Web browser engine |
| **Rendering file** | A local HTML file | HTML file on-the-fly |

### File formats

PanGraphViewer mainly uses the newly proposed reference graphical fragment assembly (rGFA) format designed in minigraph [10] as the resource to fetch and display pangenome subgraphs. However, it may not be possible to obtain high-quality genome assemblies for different individuals to generate an rGFA file using minigraph. To resolve this problem, alternative solutions have been provided in PanGraphViewer to accept the GFA_v1 [15] and VCF [16] files. Basically, the input GFA_v1 or VCF file is first converted to an rGFA file and then loaded for rendering in PanGraphViewer.

As most genome graph viewers do not offer a function to link graph nodes with genome annotations, in PanGraphViewer, we enabled this function by accepting the commonly used genome annotation formats, for example, the browser extensible data (BED) format and the general transfer/feature format (GTF/GFF) [17]. By reading the information in the corresponding genome annotation files, PanGraphViewer helps to identify gene models that are interrupted by insertions or deletions in particular samples.

### Graph visualization

In contrast to the commonly used genome graph viewers that render the entire graphs for all chromosomes or render subgraphs by nodes IDs, PanGraphViewer provides a subgraph visualization function by specifying particular chromosomal regions using coordinates or node IDs. Specifically, PanGraphViewer first fetches the subgraph information from the entire pangenome graph file and then parses and saves the requested information, for instance, chromosome ID, node and connection information into an HTML file on disk in the desktop-based version or generates on-the-fly in the web-based version. By invoking the rendering engine implemented in either PyQt5 or a web browser, subgraphs are displayed on the HTML5 canvas where users can use a mouse to drag and zoom in and out of the graph to browse the nodes of interest. By hovering on the nodes, users may read the information underlying, for example, the node ID, corresponding sample name and the sequence length.

To suit the host system and optimize performance, the desktop-based PanGraphViewer was designed to use vis.js (https://visjs.org) and cytoscape.js [18] to render graphs, while the web-based PanGraphViewer was designed to only use cytoscape.js. One of the reasons leading to this design is that cytoscape.js renders faster than vis.js when there are thousands of nodes and the computing resource consumed is less than that in vis.js. In consideration of response time, we only enable cytoscape.js for the web-based application whereas, in the desktop-based PanGraphViewer, it is allowed to switch the two JavaScript libraries by specifying the maximum number of nodes plotted in Settings to meet individual preferences. Due to the limitations of JavaScript libraries, PanGraphViewer is configured to render up to 20,000 nodes in one plot. Although it is possible to modify the config file to lessen the constraint, it is recommended to keep this setting as PanGraphViewer is designed to better fit for subgraph visualization than the whole graph visualization in consideration of the overall performance and computing resource used.

### Benchmarking

To test the stability of PanGraphViewer, we performed beta tests on Windows 10, Linux Ubuntu 20.04, Linux Ubuntu 18.04, macOS Big Sur and macOS Monterey (Intel and M1) operating systems. Meanwhile, the GitHub repository has been publicly available for free access. To test the memory and time consumed by PanGraphViewer in processing different pangenome graph files, we compared the performance of PanGraphViewer v1.0.2, Bandage v0.8.1 [11], BandageNG v2022.06 (https://github.com/asl/BandageNG) and GfaViz v0.1 Alpha [12]. The main reasons to select those tools are that all of them can accept the rGFA format and they all have a desktop-based version. As the desktop-based PanGraphViewer and the web-based PanGraphViewer share most of the codes except codes for the user interface and graph rendering, the benchmarking for the web-based PanGraphViewer was also taken by using Google Chrome v104.0.5112.81.

The benchmarking was performed on a Microsoft windows operating system (64-bit Windows 10 Enterprise platform with an Intel^®^ Core^™^ i7-9700K CPU at 3.60 GHz and 32 GB of RAM) in consideration of the big market share of the Windows system. During benchmarking, memory usage information was extracted from Task Manager installed in the Windows operating system. A stopwatch was used to measure the time taken from program launching, file loading to graph rendering.

The pangenome graph files used for benchmarking were generated from different species. Specifically, the pangenome graphs were firstly constructed using minigraph [10] on a Linux server, with three human genome assemblies (GRCh38, hg19, and T2T-CHM13 [19]), six *Listeria monocytogenes* genome assemblies [20], eight *Arabidopsis* genome assemblies [21, 22], 33 rice genome assemblies [23], 27 soybean genome assemblies [5, 24], and 26 maize genome assemblies [25] respectively. The settings for minigraph were the default ones and the backbone references for the test pangenome graphs were hg38 (human), AE017262 (*L. monocytogenes*), Col-CEN (*Arabidopsis*), IRGSP-1.0 (rice), Wm82 v4 (soybean), and B73 (maize).

During benchmarking, all pangenome graph files were imported to the selected tools. As PanGraphViewer was not specifically designed to plot graphs from all chromosomes and there is a limitation on the number of nodes in one PanGraphViewer plot, it is difficult to perform the benchmarking for a plot with more than 20,000 nodes. To solve this problem, we selected a series of subgraph regions on all pangenome graphs with less than 20,000 nodes to benchmark. Because Bandage and BandageNG cannot plot subgraphs using coordinates but node IDs, we used gfatools v0.5 (https://github.com/lh3/gfatools) to obtain the node IDs from the target subgraph regions and then benchmarked the execution time and computing resource used to render the selected subgraphs.

## Results

We have developed PanGraphViewer to assist pangenome graph visualization. Different from other genome graph viewers (**Table 2**), PanGraphViewer provides both a desktop-based version and a web-based version to meet users’ preferences. The data format accepted in PanGraphViewer is not restrained to one format. The support of subgraph visualization using coordinates enables an easy graph viewing from the entire pangenome graph without the pre-knowledge of particular nodes. Detailed node information, such as the node origin and nodes related genomic variations can be easily browsed through mouseover in the graph. The node sizes are logarithmic proportional to the number of sequences; the node shapes reflect the variant types, and the colors represent nodes for different individuals. The association with genome annotation enable users to browse gene models that are interrupted in particular samples by insertions or deletions.

**Table 2.** Comparison between PanGraphViewer and other graph viewers.

| Item | PanGraphViewer | Bandage | BandageNG | GfaViz | Sequence Tube Map | MoMI-G |
| --- | --- | --- | --- | --- | --- | --- |
| Pangenome graph | Assembly-based;<br>Variation-based | Assembly-based | Assembly-based | Assembly-based | Variation-based | Variation-based |
| Pangenome graph format | rGFA; GFA_v1; VCF | GFA_v1; rGFA | GFA_v1; rGFA | GFA_v1; rGFA | xg+gbwt+gam | xg+pcf |
| Back end | cytoscape.js; viz.js | Open Graph Drawing Framework | Open Graph Drawing Framework | Open Graph Drawing Framework | Not applicable | Sequence Tube Map; Circos |
| Version | Desktop-based;<br>Web-based | Desktop-based | Desktop-based | Desktop-based | Web-based | Web-based |
| View-level | Segment-level;<br>Chromosome-level | Chromosome-level; Segment-level | Chromosome-level; Segment-level | Chromosome-level | Nucleotide base-level; Segment-level; Chromosome-level; | Nucleotide base-level; Segment-level; Chromosome-level; |
| Target graph | Subgraph | Entire graph | Entire graph | Entire graph | Subgraph | Subgraph |
| Subgraph view | By coordinates; up to 20,000 nodes | By node IDs (node list string limit to 32,767) | By node IDs (node list string limit to 32,767) | By node IDs | By path | By path |
| Annotation format | BED, GTF; GFF3 | None | BED (under development) | None | BED | BED |

### Functions implemented in PanGraphViewer

PanGraphViewer provides different functions to facilitate pangenome graph visualization (**Figure 1**). Initially, PanGraphViewer loads a pangenome graph file in the rGFA, GFA_v1, or VCF format. In the rGFA and VCF formats, sample names or identifiers are recommended to be included in the file tags, such as ‘SN’ in rGFA to be read and shown in the PanGraphViewer plot, whereas in the GFA_v1 file, paths (‘P’) are required. The use of a VCF file may be convenient for users to visualize different variants, such as single nucleotide polymorphisms, indels, and structural variants, among individuals in a pangenome study. The node shapes used in PanGraphViewer represent different types of variants.

**Figure 1.**
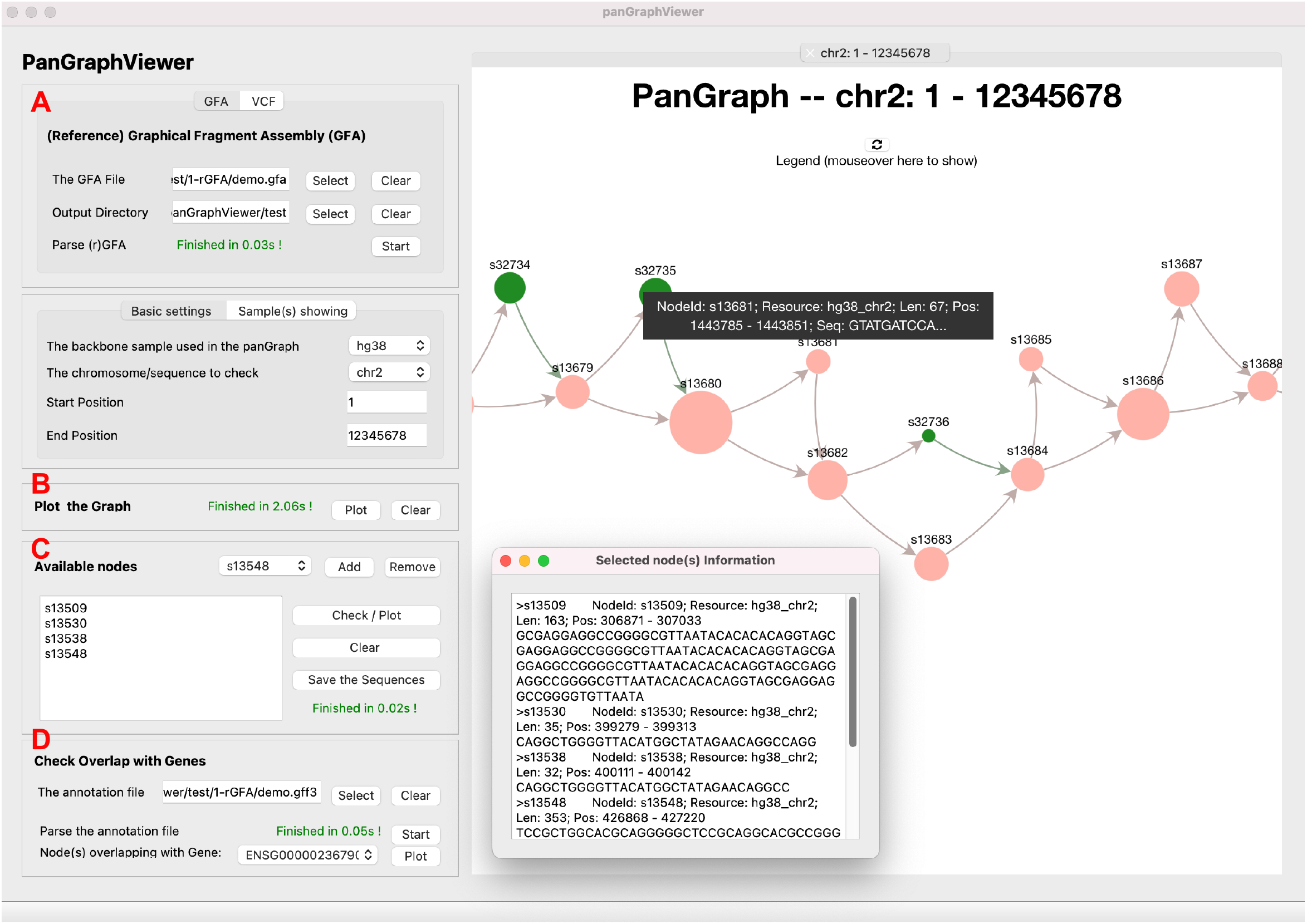
An overall view of PanGraphViewer (desktop-based version) There are four main sections in PanGraphViewer: (**A**) loading a (reference) graphical fragment assembly (GFA) file or a variant call format (VCF) file, **(B)**parsing and plotting a graph of interest, **(C)** browsing node information or plotting subgraph by node IDs, and **(D)** plotting a gene model that may be interrupted by some nodes in some samples.

When a specific chromosomal region is selected, PanGraphViewer fetches the subgraph information from the hash table created during pangenome graph file importing. The region can be any region in the pangenome. After calculating the number of nodes in the target pangenomic region and assessing, in the desktop-based PanGraphViewer, it creates an HTML file to the output directory and then loads the HTML file to the user interface canvas to be rendered by the QtWebEngine. While in the web-based PanGraphViewer, all information is processed in RAM to speed up the response time.

PanGraphViewer provides a function to remove or retrieve nodes from specific samples depending on whether the sample is of interest to the user. As pangenomic regions with variants may be of more interest, users may view or export the sequences of nodes residing in these regions for additional analyses, such as searching for homologs in a database. The DNA sequence view and export functions are available for all nodes. Furthermore, PanGraphViewer provides a function to associate genes with nodes. Through this function, users can visualize gene models that are interrupted by particular nodes in some samples due to insertions or deletions.

### Computational resources used to process different pangenome graphs

Bandage and GfaViz are commonly used to visualize genome graphs with a GFA file or an rGFA file as the input. To investigate the computational resource usage when processing different pangenome graphs, we compared PanGraphViewer, Bandage, BandageNG (the forked Bandage) and GfaViz in consideration of the accepted data format and the desktop-based version that they have. Due to the design difference, for example, Bandage and BandageNG cannot directly plot subgraphs from the entire graph using coordinates. Instead, they require users to specific the nodes with node IDs. PanGraphViewer cannot effectively plot a graph with more than 20,000 nodes. It is difficult to benchmark the performance at the same level. To minimize the effect and display the feature of pangenome graphs in a region, we selected a series of coordinates in all test pangenome graph files and employed a third-party tool (gfatools) to extract the node IDs and then plotted the subgraphs underlying. In PanGraphViewer, to make the benchmarking at the same level, we have enabled the plot function by node IDs.

During the benchmarking, GfaViz failed to execute the *Arabidopsis,* rice, soybean, and maize (the most complicated in the test) pangenome graphs under the test environment (32 Gb of RAM) (**Figure 2, supplementary Table S1-S5**), suggesting that GfaViz is not suitable for visualizing pangenome graphs with many nodes and edges on low-configuration machines. The reason leading to this failure might be that GfaViz needs to load, parse and render the entire graph from all chromosomes in one execution and this design strategy needs a large amount of time and RAM to process a big pangenome graph. Bandage, BandageNG (the forked Bandage) and PanGraphViewer, however, could load all test pangenome graph files under the test environment.

**Figure 2.**
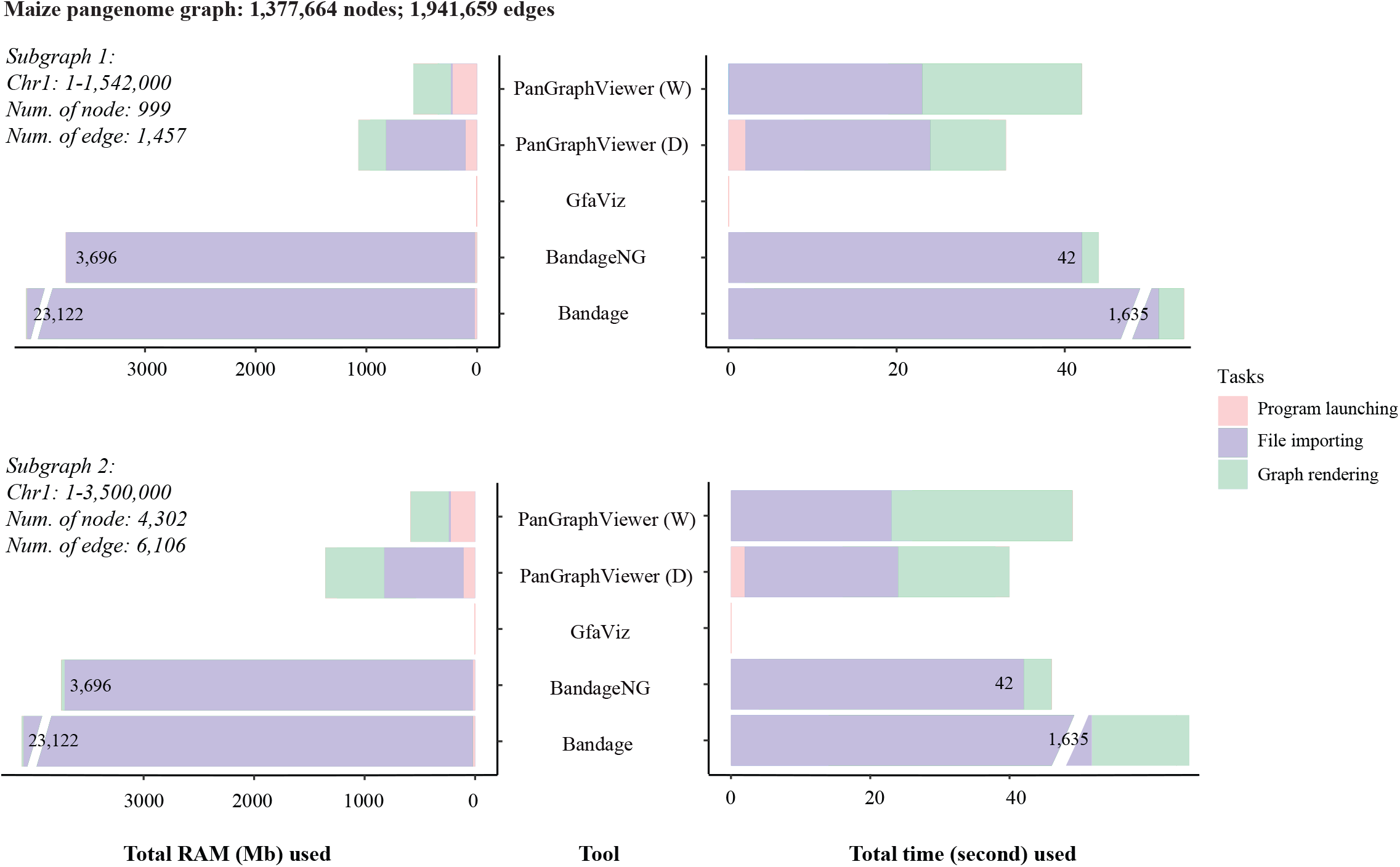
RAM and time used to render subgraphs from a pangenome graph using node ID. PanGraphViewer v1.0.2, Bandage v0.8.1, Bandage NG v2022.06 (the forked Bandage) and GfaViz v0.1 Alpha were used to benchmark the execution time and peak RAM usage when loading a pangenome graph file and then drawing subgraphs from the entire pangenome. The demo pangenome graph file (the most complicated one during the benchmarking) was generated from 26 maize genome assemblies with B37 as the backbone. Subgraphs from two pangenomic regions (Chr1:1-1,542,000 and Chr1:1-3,500,000) were selected to be viewed through the application of node IDs. During pangenome graph file loading, PanGraphViewer (D: desktop-based version; W: Web-based version) consumed the least amount of RAM and time, while GfaViz failed to import and render graphs from the maize pangenome due to insufficient memory (32 GB) in the test machine. The performance of BandageNG was better than Bandage. However, the execution time and computing resources used in BandgeNG were still higher than that used in PanGraphViewer. This might be that during design, the two Bandage tools need to process the entire pangenome graph file into memory, while PanGraphViewer only extract the needed information without pre-processing them. The RAM consumption of the two Bandage tools may vary depending on the size and complexity of the pangenome graph.

In most cases, the two Bandage tools required more memory than PanGraphViewer (**Figure 2, supplementary Table S1-S5**). This might be that PanGraphViewer avoids storing the entire pangenome graph information into memory while Bandage and BandageNG do. When processing the *L. monocytogenes* pangenome graph (**supplementary Table S5**), Bandage and BandageNG used much less memory than PanGraphViewer, in which the graph is much simpler than the others. This can be explained by the fact that Bandage and BandageNG use the more efficient (in both storage and speed) C++ language, which PanGraphViewer use less efficient (but more portable) Python scripts.

For execution time, Bandage used more time to import the pangenome graph files than PanGraphViewer, particularly for those with many nodes and edges (**Figure 2, supplementary Table S1-S5**). This is likely because during file loading, PanGraphViewer only reads the pangenome graph files and extracts the node ID, chromosome ID and sample sequence information. There is no further information processing during pangenome graph file loading in PanGraphViewer, while Bandage needs to perform extra analyses for example calculating the number of nodes and edges and storing the entire graph information into memory, which consumes additional time. In the forked Bandage (BandageNG), it seems the performance has been improved remarkably, though for large pangenome graphs, it also needs time to import the files (**Figure 2, supplementary Table S1-S5**).

For subgraph rendering, PanGraphViewer used more time than the two Bandage tools. This is because Bandage and BandageNG have already stored the node information in RAM during file loading and could fetch the needed information directly to generate the subgraph through the open graph drawing framework. Because of this design, the execution time could be saved but more memories are needed to pre-store the node information. In PanGraphViewer, however, it does not pre-store the node sequence information during file loading. During plotting, PanGraphViewer revisits the pangenome graph file and extracts the needed information into a HTML file on disk (desktop-based version) or on-the-fly (web-based version) and then renders the underlying subgraph. As a rendering engine is needed to display the graph, extra time are taken in PanGraphViewer. Overall, PanGraphViewer sacrifices some performance to balance the execution time and memory consumption to make it better fit for a relatively low configure machine.

Noteworthily, during the benchmarking, the time used to extract node IDs from the third-part tool was not included. As there is a string length limit (32,767) in the ‘Node(s)’ box in Bandage and BandgeNG, we could not use a larger number of nodes and edges to test. Overall, the benchmarking can roughly present the performance of the tools.

## Discussion

PanGraphViewer was designed for pangenome graph visualization. It clearly shows the shared nodes among all individuals and the variable nodes present in some individuals with different representative colours. The recommendation of pre-inputting sample names makes it possible to visualize the node origin information by hovering the mouse on the graph, which is a feature that is absent from most pangenome graph viewers. As users may have different technical backgrounds and computational resources, we prepared a desktop-based PanGraphViewer and a web-based PanGraphViewer for users to select.

The data formats accepted by graph viewers differ with some, such as Sequence Tube Map [13] and MoMI-G [14], using customized data formats. Using a customized data format may prevent data sharing and make the application of the tool complicated. To avoid such a problem, PanGraphViewer uses the rGFA format, a subset of the generic GFA format, as the main data resource. There are two reasons for using such a format. First, customized information could be easily added to the tags in an rGFA file to enrich the content (contrary to generic GFA files), without affecting the display of the graphs in other tools, such as Bandage and GfaViz. Second, in contrast to the rGFA format, the generic GFA format was not initially designed for pangenome graphs and the expansibility of the generic GFA format is limited. For instance, it is non-trivial to add the sequence coordinate and node origin information in the commonly used GFA_v1 format.

In addition to accept GFA_v1 and rGFA as the input, PanGraphViewer accepts VCF files, which are relatively easy to obtain using SV callers such as sniffles [26], to allow the integration of SV information into pangenome graph. The visualization of a pangenome graph from a VCF file may be of interest to researchers who are studying genetic variation in a species. The connection between genome annotation and graph nodes may be useful for users to identify the presence and absence of variation in a pangenome due to insertions and deletions through a plot after selecting a particular gene of interest in PanGraphViewer.

Previous pangenome graph viewers cannot directly plot subgraphs from a complete pangenome graph using coordinates. Users may need to render the entire graph for all chromosomes and then locate and browse the subgraphs of interest. For graphs with fewer nodes and connections, this strategy is efficient. However, for a pangenome graph with many chromosomes, nodes, and connections, such a solution may not be optimal, as it requires a large number of computational resources to complete the process, and users may need a machine with a relatively advanced hardware configuration. Alternatively, users could find a way to get the node IDs within a pangenomic region and then plot the underlying subgraph or they have to truncate the pangenome graph files and load the manipulated file to view. These practices may increase the difficulty of using the graph viewer and a third-party tool is also required. As the alternative approaches could only create static subgraphs, the extraction of node IDs or generation of truncated pangenome graph files would need to be taken again for another subgraph viewing. This could complicate the application of the tool.

In contrast to the commonly used graph viewers, PanGraphViewer uses a different design strategy. For example, it enables subgraph visualization by coordinates by writing the information to an HTML file (desktop-based version) or generating on-the-fly (web-based version) and then renders the graph. As the size of a pangenome graph depends on the number of individuals, PanGraphVeiwer also allows the selection of individuals. Writing subgraph information to an HTML file may enable users to further explore the subgraph using a web browser without the need to reopen the tool and render the graph. This may save time to avoid regenerating the subgraph and allow users to visualize the target pangenome graph more efficiently. Additionally, visualizing subgraphs directly by coordinates could overcome the pitfall of manually locating the target chromosome and then locating the genomic region of interest, because such a practice is not trivial, particularly when there are many chromosomes or the chromosome of interest is long. Users also don’t need to rely on a third-party tool to extract the node information and browse the underlying subgraph. This could lower the difficulty in viewing a pangenome graph and users can easily check the variations in a particular pangenomic region to study the biological meaning underlying.

PanGraphViewer has some limitations. For example, it cannot efficiently render a graph with more than 20,000 nodes due to the limitations inherited from the JavaScript libraries that are used. The layout of the entire pangenome graph from different chromosomes may not be able to be shown in PanGraphViewer for large pangenomes. However, as subgraph viewing is the main target for PanGraphViewer, 20,000 nodes might be sufficient for users to explore. Notably, the subgraph visualization function provided in PanGraphViewer enables users to easily browse specific genome regions using coordinates with few computational resources used, whereas in other tools more RAM and time may be needed to achieve this. Furthermore, the overall structure of the entire pangenome graph from other tools may have a low resolution particularly if the graph is big, users may get limited information in this case. Detailed information in a subgraph region may be more interesting to researchers in a pangenome study. Admittedly, subgraph viewing has its weakness. For instance, if users have no idea on which coordinates to target for, they cannot benefit from subgraph viewing at the beginning. Users may need to view and explore a larger region or even a chromosome first, and then input the target regions after exploration. PanGraphViewer, however, can help to achieve this goal.

## Conclusions

PanGraphViewer is versatile and efficient for pangenome graph visualization, irrespective of the operating system. It is easy to install and deploy. The functions provided in PanGraphViewer may be of particular interest to researchers performing pangenome analyses, for instance, visualizing different structural variations in the studied samples. Given the absence of different pangenome graph viewers, we expect PanGraphViewer to be a useful tool for pangenome studies. Further development will be conducted to expand the current pangenome graph visualization from a species level to a genus level using for example genes as the backbone. Haplotype structure visualization in a pangenome would also be included.

## Data availability

The pangenome graph files constructed in the study are at https://doi.org/10.6084/m9.figshare.20296575.v1

## Code availability

PanGraphViewer is freely available at https://github.com/TF-Chan-Lab/panGraphViewer, licensed under the MIT licence.

## CRediT author statement

**Yuxuan Yuan**: Conceptualization, Software, Formal analysis, Writing - original draft, Writing - review & editing. **Ricky Ka-Kui Ma**: Software, Writing - review & editing. **Ting-Fung Chan**: Conceptualization, Writing - review & editing, Supervision. All authors read and approved the final version of the manuscript.

## Acknowledgements

This work was supported by the Hong Kong Research Grants Council Area of Excellence Scheme (AoE/M-403/16) and Collaborative Research Fund (C4057-18EF) and the Innovation and Technology Commission, Hong Kong Special Administrative Region Government to the State Key Laboratory of Agrobiotechnology (CUHK). Any opinions, findings, conclusions, or recommendations expressed in this publication do not reflect the views of the Government of the Hong Kong Special Administrative Region or the Innovation and Technology Commission.

